# Thalamic Dysconnectivity and Negative Symptoms in Schizophrenia Spectrum Disorder

**DOI:** 10.64898/2026.10.04.756569

**Authors:** Keiko Kunitoki, Bhim M. Adhikari, A. Ankeeta, Si Gao, Yizhou Ma, Kelsey Straub, Gustavo Dos Santos Alves Maria, Antonio F. Pagán, Alia Warner, Xiaoming Du, Peter Kochunov, L Elliot Hong

## Abstract

**Background and Hypothesis:** Negative symptoms in schizophrenia spectrum disorder (SSD) are associated with poor functional outcomes and reduced quality of life; however, effective treatments remain limited. A better understanding of the underlying brain mechanisms is needed to develop better treatments. Neuroimaging studies implicate multiple brain networks in negative symptoms, though findings remain heterogeneous, potentially reflecting the multidimensionality of both symptoms and brain networks. Larger, more systematic dissections are required to identify the key hub(s) underlying negative symptoms.

**Study Design:** Structural MRI data from 394 participants (210 SSD, 184 healthy controls [HC]) were analyzed to identify cortical thickness and subcortical volume demonstrating both SSD-HC differences and associations with negative symptoms measured by the Brief Negative Symptom Scale (BNSS). Regions meeting both criteria were selected as seeds for resting-state functional connectivity analyses in 143 SSD patients to examine association with negative symptoms and their subdomains.

**Study Results:** The thalamus was the only region demonstrating significant structural abnormalities both in SSD and an association with BNSS total score (p-FDR<0.05). Functional-network-defined thalamic subregions were subsequently examined. Several thalamic hypoconnectivities were significantly corresponded with negative symptoms, involving thalamic subregions associated with the cortical default mode, somatosensory, medial temporal, and dorsal attention networks (p=0.001, *a*_corrected_<0.05). Blunted affect showed the most extensive thalamic connectivity alterations, involving the striatum, cerebellum, and temporoparietal junction/superior temporal sulcus (TPJ- STS). Thalamic hypoconnectivity with TPJ-STS was also associated with overall negative symptoms and anhedonia.

**Conclusions:** This study identifies the thalamus as a central hub of negative symptom-related dysfunction in SSD and highlights specific thalamic subnetworks as potential targets for circuit-based interventions.

Keyword: Schizophrenia, Negative Symptoms, Structural MRI, Functional Connectivity, Thalamus

## Introduction

Negative symptoms in schizophrenia spectrum disorder (SSD) refer to a symptom cluster of absence or reduction of motivation, social drive, emotion and emotional expression, speech, and other activities. SSD patients with more severe negative symptoms have poorer functioning, recovery-based outcomes, and quality of life.^1^ Despite their clinical significance, treatment of negative symptoms remains difficult, and most patients with negative symptoms show no or partial response to current pharmacological and psychosocial interventions.^2,3^ Among antipsychotics, cariprazine,^4^ clozapine,^5^ and xanomeline^6^ have demonstrated relatively better but modest reduction in negative symptoms and may partially be attributed to secondary improvements from reductions in positive symptom^7^. Other adjunct treatment options include antidepressants^8,9^, antibiotics such as minocycline, immunomodulators, psychosocial interventions, and physical activity, although study quality in many of these trials is low to moderate^10^ and none of them are currently considered as gold-standard treatments of negative symptoms^3,7,10–13^. Consequently, identifying the neurobiological mechanisms underlying negative symptoms is critical to guide the development of more effective, targeted interventions.

Large-scale structural neuroimaging studies from the ENIGMA consortium have identified widespread cortical thinning in SSD, particularly within frontal and temporal regions ^14^, as well as reduced volumes of the hippocampus, amygdala, thalamus, and nucleus accumbens ^15^. More recently, meta-analytic evidence has highlighted the thalamus and amygdala as among the most consistently reduced subcortical structures in individuals with SSD compared with healthy controls ^16^. It is likely some of these structural abnormalities may be involved in negative symptoms. Indeed, studies focused on individuals with prominent negative symptoms have linked persistent negative symptoms to structural alterations across distributed cortical and subcortical regions, including the prefrontal cortex, frontoparietal networks, anterior cingulate cortex, insula, cerebellum, and several subcortical structures including thalamus and amygdala ^14,17–21^. Functional neuroimaging studies likewise suggest that negative symptoms in SSD arise from dysfunction within distributed neural circuits and their constituent nodes, rather than from isolated regional abnormalities.^22,23^ Implicated regions include the cerebellum,^24–27^, precuneus^28–30^, insula^26,30,31^, thalamus^21,31^, ventral tegmental area/substantia nigra^32,33^, striatum ^34,35^, dorsolateral prefrontal cortex ^36^, medial orbitofrontal cortex ^37^, and amygdala^38^. Consistent with these findings, functional connectivity studies have further implicated multiple large-scale brain networks in the pathophysiology of negative symptoms, including the default mode, salience, frontoparietal control, sensorimotor, affective, auditory-language, and subcortical networks^22,23,39–49^. However, findings are quite heterogeneous and often inconsistent, which may suggest certain underlying dysfunction that engages multiple networks and different patients or cohorts may be associated with only part of these abnormalities. Alternatively, it may result from the heterogeneity of negative symptoms themselves and inconsistencies in their measurement, which would require more fine-grained analysis of brain circuits associated with specific subcomponents of negative symptoms.

Another important challenge in identifying the brain circuits associated with negative symptoms is that SSD involves widespread abnormalities across multiple brain regions and neural circuits, whereas only a subset of these abnormalities is specifically linked to negative symptoms. We propose to first use structural imaging data to identify brain regions that are generally affected in SSD patients and with negative symptoms, subsequently utilize the structural abnormalities to guide the search for functional circuits most likely contributing to negative symptoms, aiming to identify the brain regions/circuits more likely contributing to negative symptoms in SSD. Furthermore, we also hypothesized that distinct negative symptom subdomains would exhibit specific connectivity patterns, reflecting the heterogeneity of negative symptom manifestations and their underlying neural substrates.

Finally, given the associations between thalamic volume reduction and negative symptoms, and thalamus is known to consist of partially segregated regions associated with different major functional networks across the brain ^50–54^, we further subdivided the thalamus into functional-network-defined subregions. We then explored whether negative symptoms are associated with altered thalamic subregion-based functional connectivities. Identifying these symptom-specific circuits may advance mechanistic understanding of negative symptoms and support the development of circuit-based interventions targeting clinically meaningful dimensions of negative symptoms.

## Methods

### Participants

Patients with schizophrenia spectrum disorder were recruited from the Maryland Psychiatric Research Center and mental health clinics in the greater Baltimore area. HC were recruited through media advertisements and word of mouth from the same geographic area. The Structured Clinical Interview for DSM-IV Axis I Disorders (SCID-I) or DSM-V was used to confirm diagnosis or the lack thereof for all participants. History of major medical and neurological disorders, head injury with cognitive sequelae, intellectual disability, and substance abuse or dependence within the past 6 months (except nicotine and cannabis) were exclusionary. Individuals with past or present DSM-IV Axis I psychiatric disorders or family history of psychosis in the prior two generations were excluded from healthy controls. Of the 210 patients, 200 reported medication information, including 25 who were not taking antipsychotic medication at the time of the study. The most frequently used antipsychotics were clozapine and risperidone (used by 40 patients each) and aripiprazole (used by 30 patients). Chlorpromazine equivalents (CPZ) were calculated based on the International Consensus Study of Antipsychotic Dosing (ICSAD)-2, or when not available, ICSAD-1.^55,56^ In 39 patients, partial information for frequency or daily dose were missing or could not be confirmed; if medication frequency was missing, it was assumed to be a daily dose; if dose information was missing, the midpoint of the target dose range defined by ICSAD was used. We removed two participants who reported a daily CPZ equivalent dose of 4,000 mg or more after confirming clinically unreasonable dosing. Written informed consent was obtained from all participants as approved by the University of Maryland IRB.

We used the Brief Negative Symptoms Scale (BNSS)^57^ to measure negative symptoms. The BNSS is a 13-item clinician-rated instrument, designed to measure 6 subscales of negative symptoms in schizophrenia: anhedonia, avolition, asociality, blunted affect, alogia, and lack of normal distress. We also used the Brief Psychiatric Rating Scale (BPRS) total, positive symptoms subscale (unusual thought content, hallucinatory behavior, conceptual disorganization, and grandiosity), and negative symptoms subscale (motor retardation, blunted affect, and emotional withdrawal) scores^58^. For cognitive function measurement, we used the Digit Symbol Substitution test (for processing speed) from the Wechsler Adult Intelligence Scale-3^59^ [26] and Digit Sequencing task (for working memory) from the Brief Assessment of Cognition in Schizophrenia^60^.

### MRI acquisition, processing, and data analysis

#### Structural data

MRI data were acquired at the University of Maryland Center for Brain Imaging Research using Siemens 3 T TRIO MRI or PRISMA (Erlangen, Germany) system. High-resolution T_1_-weighted structural images were acquired using a fast spoiled gradient-recalled echo sequence. For the Siemens TRIO scanner with 32-channel phase array head coil, acquisition parameters were: repetition time (TR) = 2200 ms, echo time (TE) = 2.83 ms, flip angle = 13°, matrix size = 220 × 320, 208 sagittal slices, and voxel size = 0.8 mm isotropic. For the Siemens PRISMA scanner with 64-channel phase array head coil, acquisition parameters were: TR = 2400 ms, TE = 2.22 ms, flip angle = 8°, matrix size = 300 × 320, 208 sagittal slices, and voxel size = 0.8 mm isotropic. Structural images were processed using FreeSurfer ‘*recon-all’* pipeline ^61^, which includes skull stripping, surface reconstruction, and quality control on cortical surfaces, subcortical segmentation, and image alignment. Cortical thickness measures were extracted based on the Desikan-Killiany atlas ^62^, yielding 34 regions per hemisphere. Subcortical volumes, including the thalamus, caudate, putamen, pallidum, hippocampus, amygdala, and nucleus accumbens, were derived using the automated segmentation Aseg atlas ^63^. Imaging protocols were adjusted from the TRIO scanner to the PRISMA scanner. For structural data, N=273 were from TRIO and N=121 from PRISMA. Structural features extracted, from the two scanner systems, were processed with the parametric implementation of NeuroCombat ^64^. NeuroCombat removes scanner effects by estimating scanner-specific location and scale parameters for each imaging feature and applying an empirical Bayes framework that pools information across features to improve parameter estimation. Cortical thickness and subcortical volume were harmonized in separate NeuroCombat models. Age and sex were included as biological covariates to preserve variance associated with these factors during harmonization. For statistical analysis, we used average value from bilateral hemispheres for each ROI. We conducted separate linear regression analyses to examine group differences in each of the 41 structural regions of interest (ROIs) demonstrating differences between individuals with SSD and HC. All structural MRI analyses were conducted in R (version 4.5.1). Age, sex, and intracranial volume were included as covariates in all models. False discovery rate (FDR)-corrected p-values < 0.05 were considered statistically significant, with correction applied across all ROI-level tests.

#### Resting state functional MRI data

Resting-state functional T_2_*-weighted images were acquired with the following imaging parameters; TRIO: repetition time (TR) = 2000 ms, echo time (TE) = 27 ms, flip angle = 90°, matrix size = 128 × 128, 37 slices, voxel size = 1.72 × 1.72 × 4 mm³, and 444 volumes. PRISMA: TR = 780 ms, TE = 34 ms, flip angle = 52°, matrix size = 104 × 104, 72 slices, voxel size = 2 mm isotropic, and 420 volumes, using an interleaved multiband acceleration factor of 8. In addition, anterior-to-posterior and posterior-to-anterior phase-encoded images were acquired for each participant to facilitate correction of susceptibility-induced distortions. In PRISMA scanner, participants underwent rsfMRI data acquisition that consisted of two runs. Oblique axial acquisitions alternated between phase encoding in the anterior-to-posterior (AP) and posterior-to-anterior (PA) directions within a single run. Separate single-band reference images, acquired for phase encoding in AP and PA directions, were used for spatial distortion correction. During the scan, participants were asked to keep their eyes closed and relax. The ENIGMA resting-state analysis pipeline implemented in the Analysis of Functional NeuroImages (AFNI) software ^65^ was used to process the resting-state fMRI (rsfMRI) data. This single-modality resting-state analysis pipeline is an extension of the conventional AFNI rsfMRI pipeline ^66^. Preprocessing included PCA-based denoising to improve signal-to-noise ratio, despiking, slice-timing correction, motion correction, registration of the base volume to the ENIGMA EPI template ^67^ and smoothing using a 4 mm full-width-half-maximum Gaussian kernel. As a part of the pipeline, time points with excessive motion (>0.2 mm estimated displacement from one time point to the next), including neighboring time points and outlier voxels fraction (>0.1) were censored from statistical analysis.

To investigate detailed thalamic contributions to negative symptom dimensions, we tested seed regions based on functional thalamic atlas (https://identifiers.org/neurovault.image:111302). The functional thalamic subregions ^52^ were derived from resting-state fMRI using a winner-take-all network assignment approach. Each thalamic voxel was assigned to the cortical functional network with which it demonstrated the strongest partial functional connectivity, after controlling for connectivity with all other networks. These thalamic subregions were labelled by the thalamo-cortical functional connectivity that have identified as follows; 1: default mode, 2: salience, 3: somatomotor, 4: executive, 5: lateral occipital (visual), 6: medial occipital (visual), 7: medial temporal (part of default mode), 8: temporal (part of somatomotor), and 9: superior fronto-parietal (dorsal attention), averaged across hemispheres. In our study, the mean time series extracted from these regions were correlated with all other brain voxels time series using Pearson correlation coefficients to generate whole-brain connectivity maps separately. Correlation coefficients were subsequently transformed to Fisher’s z-scores for group-level analyses. Multiple linear regression analyses were conducted using ‘*3dttest++*’ within the SSD group to examine associations between seed-based functional connectivity and BNSS total score and also the six subscale scores, implementing the adjustment of age and sex. Significant results were thresholded using Monte Carlo simulations by AFNI’s 3dClustSim with autocorrelation function option to yield whole-brain results thresholded at corrected alpha <0.05/n, n=number of ROIs or number comparisons corresponding to cluster-size threshold of voxels at voxel-wise threshold, p=0.001; the number of voxels required for corrected α, in this study, for 9 and 54 comparisons were 89 and 129 respectively. Then, subject-specific estimates of resting-state functional connectivity (rsFC) analysis in those significant clusters were extracted. As this study is a secondary analysis of multiple original studies, not all of the participants with SSD received rs-fMRI. Among 143 rs-fMRI from SSD patients, 80 were from TRIO and the remaining 63 were from PRISMA. The connectivity measures derived from the two scanners were further processed with the parametric implementation of NeuroCombat ^64^, which has been widely applied to rsFC measures in multisite studies, including SSD cohorts^68,69^, to account for scanner-related effects ^70–72^. The resulted harmonized rsFC data were used for further analysis with age, sex, and BNSS subdomain scores as biological covariates.

#### Additional Statistical Methods

Group comparisons of demographic, clinical, and cognitive characteristics between SSD vs HC groups were evaluated using independent-samples *t*-tests for continuous variables and Pearson’s Chi-square tests for categorical variables. Correlations among CPZ equivalent dose, clinical measurements, and average thalamic volumes were calculated with multivariable linear regression models. Age, sex, and intracranial volume were adjusted for structural MRI values.

## Results

### Structural imaging findings

A total of 394 participants (210 SSD patients and 184 HC) were included in the structural brain imaging analysis; age did not significantly differ, but there were significantly more males in the SSD group (**Table 1**). The mean duration of illness was 16.4 years (SD = 14.0). For cortical thickness of the 34 cortical ROIs and subcortical gray matter volume of the 7 subcortical ROIs, we found significant group differences in 29 cortical ROIs and 3 subcortical ROIs, all due to thinner cortex or smaller subcortical volumes in SSD (**Figure 1A**). Among these ROIs, only thalamus volume was significantly correlated with negative symptoms as measured by BNSS total scores among all 205 SSD patients dataset (β=-0.18, p=0.001, p-FDR=0.04) (**Figure 1B-C**).

**Figure 1.**
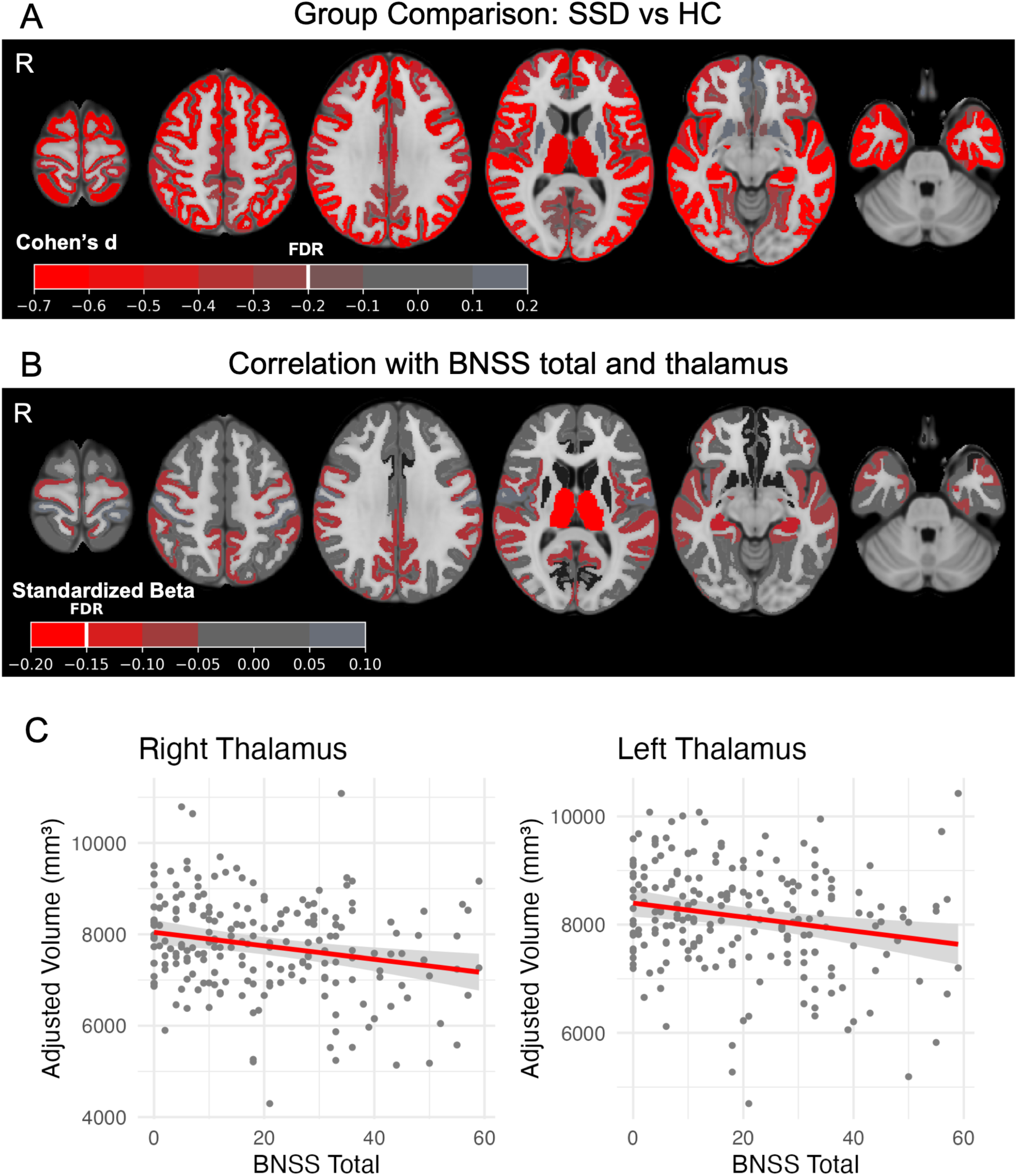
A: Effect size map of group comparison between SSD and HC. Significant group differences were found in 29 cortical regions and 3 subcortical regions including thalamus. Color coded by Cohen’s d effect size. Heavy line on the color bar indicates significance threshold after FDR correction. SSD n=210, HC n=184. B: Regional assocaitions with BNSS total score within SSD group. Average thalamus volume was negatively associated with BNSS total score. Color coded by standardized beta (ß) with heavy line in the color bar indicates theoretical threshould for significance after FDR correction. Regions not significantly different between SSD vs HC are excluded from the analysis and blacked out in the figure. SSD n=205. C: Correlation between BNSS total score and right and left thalamus volume within SSD group. Age, sex, and intracranal volume were adjusted.

**Table 1.** Demographics for structural MRI data for individuals with SSD and HC groups.

|  | **SSD**<br>N = 210 | **HC**<br>N = 184 | **Statistic** | **p-value** |
| --- | --- | --- | --- | --- |
| **Age** | 37.7 (14.0) | 38.0 (14.3) | t = -0.18 | >0.9 |
| **Gender (M/F)** | 143/67 | 91/93 | $\chi^2 = 13$ | <0.001 |
| **Race/Ethnicity** | | | $\chi^2 = 17$ | 0.005 |
| Hispanic | 7 (3.3%) | 4 (2.2%) |  |  |
| American Indian/Alaska Native | 1 (0.5%) | 0 (0%) |  |  |
| Asian | 5 (2.4%) | 16 (8.7%) |  |  |
| Black or African American | 92 (44%) | 52 (28%) |  |  |
| Multiracial | 4 (1.9%) | 4 (2.2%) |  |  |
| White | 94 (45%) | 100 (54%) |  |  |
| Unknown | 7 (3.3%) | 8 (4.3%) |  |  |
| **Current Smoker** | 75 (36%) | 45 (24%) | $\chi^2 = 5.3$ | 0.015 |
| **Duration of illness (years)** | 16.4 (14.0) | - |  |  |
| **Education** | | | $\chi^2 = 48$ | <0.001 |
| Less than high school | 34 (16%) | 5 (2.7%) |  |  |
| High school diploma or GED | 59 (28%) | 26 (14%) |  |  |
| Some college or Associate's degree | 80 (38%) | 75 (41%) |  |  |
| Bachelor's degree or higher | 37 (18%) | 78 (42%) |  |  |
| **Scanner (Trio/Prisma)** | 138/72 | 135/49 | $\chi^2 = 2.7$ | 0.10 |
| **CPZ equivalent dose** | 526.2 (431.7) | - |  |  |
| **BPRS total** | 35.8 (11.1) | - |  |  |
| Positive | 9.5 (4.8) | - |  |  |
| Negative | 6.3 (2.9) | - |  |  |
| **BNSS total** | 19.9 (15.6) | - |  |  |
| Anhedonia | 4.8 (4.8) | - |  |  |
| Distress | 0.7 (1.2) | - |  |  |
| Asociality | 3.3 (2.8) | - |  |  |
| Avolition | 3.6 (3.1) | - |  |  |
| Blunted Affect | 5.1 (4.8) | - |  |  |
| Alogia | 2.5 (3.0) | - |  |  |
| **Digit Sequencing** | 17.3 (4.9) | 21.3 (4.3) | t = -8.3 | <0.001 |
| **Digit Symbol Coding** | 57.7 (19.6) | 77.2 (17.0) | t = -10 | <0.001 |

### Resting state functional connectivity (rsFC) findings

We then examined whether rsFC of the thalamus and/or its sub-nuclei are associated with negative symptoms. One-hundred and forty-three SSD patients were included in the seed-based rsfMRI analysis with the thalamus as well as nine bilateral functional thalamic subregions as seeds (**Figure 2A**). Overlap between functional thalamic subregions and histology-based thalamic nuclei are shown in **Figure 2B**. BNSS total score was not significantly associated with whole-thalamic seed-based rsFC. BNSS total score was negatively associated with rsFC between five thalamic subregions and three other brain regions. Specifically, BNSS total was negatively associated with rsFC between three thalamic subregions (somatomotor, medial temporal, and dorsal attention) and the temporoparietal junction to superior temporal sulcus (TPJ-STS) (**Figure 2C and 2D**) as well as rsFC between two additional thalamic subregions (default mode and medial occipital) and the cerebellum (**Figure 2E**).

**Figure 2.**
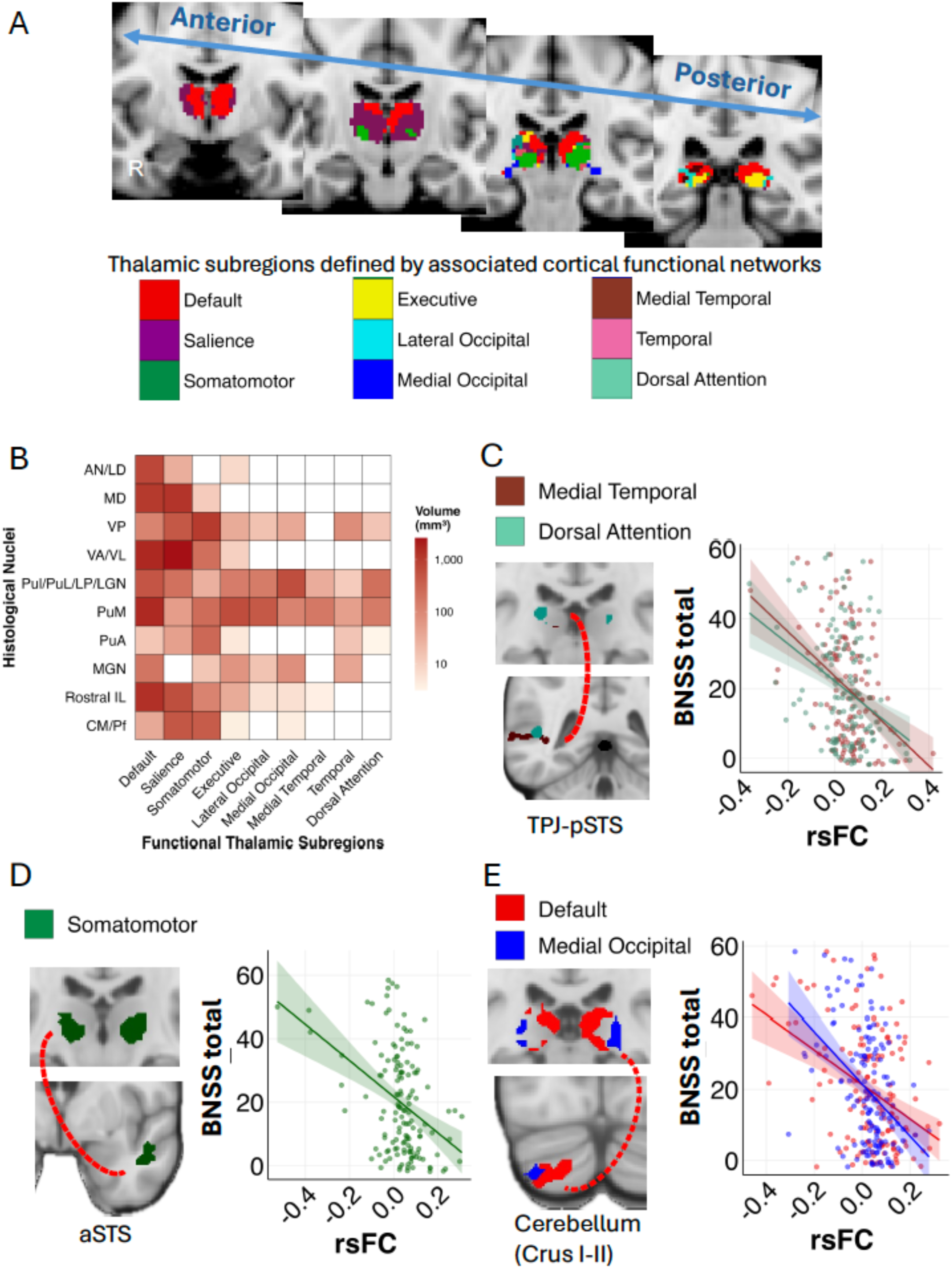
Resting-state functional connectivity associated with BNSS total score within the SSD group. A. Functional thalamic subregion atlas based on thalamo-cortical functional network. 9 subregions are color-coded and named for corresponding cortical network. B. Heat map of volumetric overlap between functional thalamic subregions and histological thalamic nuclei^134–137^. The color bar is in log10 scale. C-E. BNSS total score was associated with five connectivity measures involving five thalamic subregions and three other brain regions (n=143). These connections were between the medial temporal lobe-related and dorsal attention network-derived thalamic subregions and TPJ-pSTS (C), somatomotor network-derived thalamic subregion and aSTS (D), and default mode network-derived and medial occipital lobe-related thalamic subregions and cerebellum (E). Harmonized and age- and sex-adjusted data were used for the visualization. STS: superior temporal sulcus. aSTS: anterior superior temporal sulcus. pSTS: posterior superior temporal sulcus. TPJ: temporoparietal junction. Thalamic nuclei groups: AN/LD: anterior and dorsolateral nuclei. MD: mediodorsal nucleus. VP: ventral posterior nucleus. VP/VL: ventral anterior and ventral lateral nuclei. PuI/PuL/LP/LGM: inferior pulvinar, lateral pulvinar, lateral posterior nucleus, and lateral geniculate nucleus. PuM: medial pulvinar, PuA: anterior pulvinar. MGN: medial geniculate nucleus. Rostral IL: rostral intralaminar nuclei. CM/Pf: centromedian-parafascicular complex.

We further tested associations among the six BNSS subdomain scores with rsFC of the entire seed based rsFC using the thalamus, as well as the nine thalamic subregions. Blunted affect demonstrated the most extensive pattern of associations with thalamic rsFC, showing significant associations with 16 rsFC from the whole thalamus and 8 out of 9 thalamic subregion seed-based rsFC to TPJ-STS, cerebellum, and striatum (**Figure 3A**). Anhedonia was significantly associated with a rsFC between thalamic subregions associated with medial temporal network and TPJ-STS (**Figure 3B**). We found no significant thalamic subregion rsFC associated with the lack of normal distress, asociality, avolition, or alogia subdomain scores.

**Figure 3.**
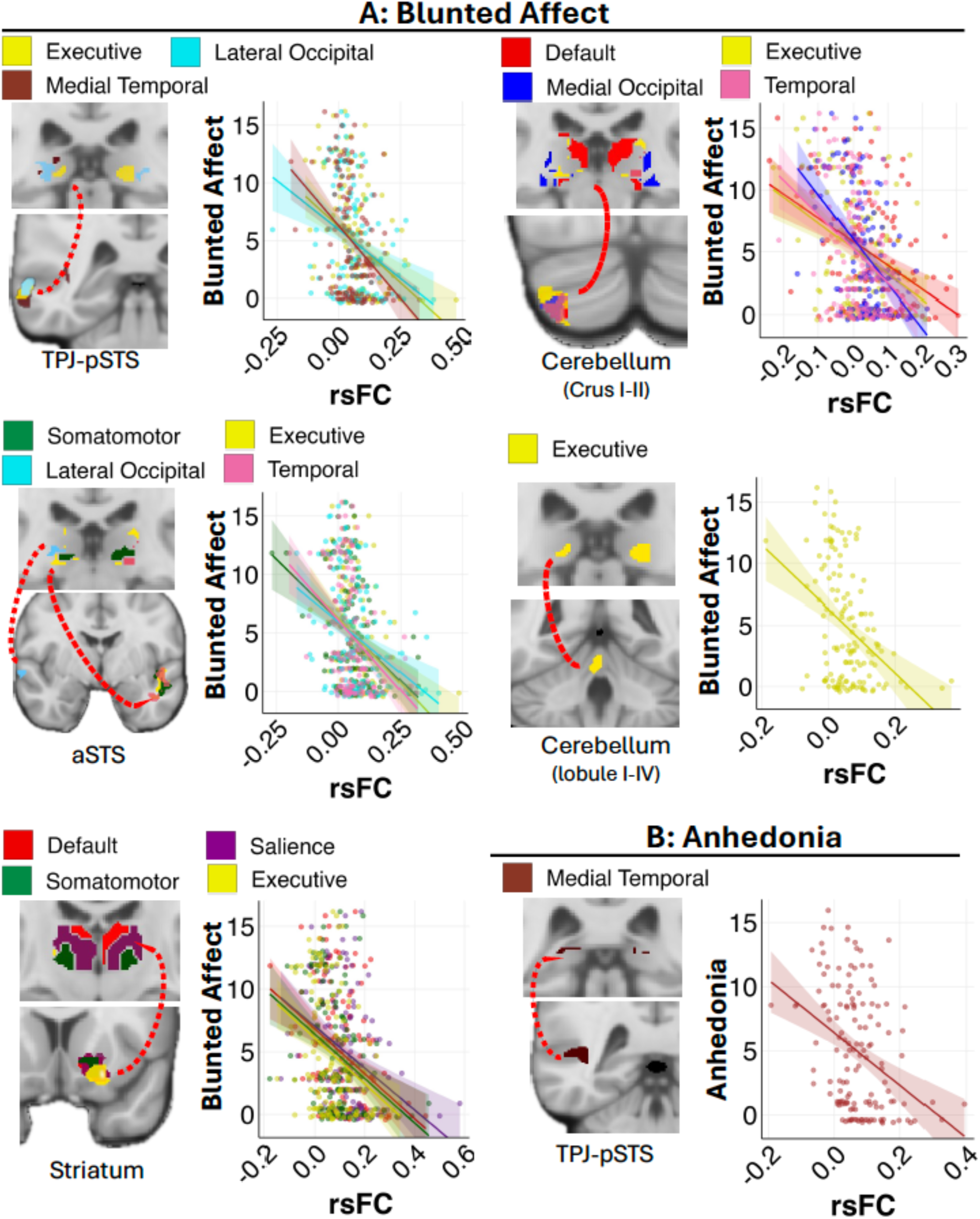
Resting-state functional connectivity associated with BNSS subscales within the SSD group. Thalamic subregions are color-coded and named based on their associated cortical lobe or functional network, as shown in Figure 2. A. Blunted affect was associated with 16 connectivity measures involving eight thalamic subregions out of nine, including connections with the temporoparietal junction, temporal lobe, cerebellum, and striatum. B. Anhedonia was associated with a connectivity measure involving the thalamic subregions associated with medial temporal network and TPJ-pSTS. Harmonized and age- and sex-adjusted data was used for the visualization. aSTS: anterior superior temporal sulcus. pSTS: posterior superior temporal sulcus. TPJ: temporoparietal junction.

### Relationships to Antipsychotics and Other Symptoms

Among SSD patients, CPZ equivalent dose was not significantly associated with either BPRS total score or BNSS total score, adjusted for age and sex (BPRS: ß=0.13, p=0.9, BNSS: ß=-0.03, p=0.7). CPZ equivalent dose was not significantly associated with bilateral thalamic volume, adjusted for age, sex, and intracranial volume (ß=-0.01, p=0.86). CPZ equivalent dose was also not significantly associated with the rsFC correlated with BNSS domains after multiple comparison correction. BNSS total score demonstrated a weak, positive association with BPRS total score (ß=0.18, p=0.013), a moderately strong, positive association with the BPRS negative symptoms subscale (ß=0.65, p<0.001),and no significant association with BPRS positive symptoms subscale (ß=-0.02, p=0.81). BNSS total score was negatively associated with processing speed (ß=-0.20, p<0.001) but not significantly associated with working memory (ß=- 0.08, p=0.26).

## Discussion

In this study, we examined the neural underpinnings of negative symptoms using structural findings and detailed rsFC analysis. We found that smaller bilateral thalamic volume was the strongest SSD-related structural deficit linked to BNSS-measured negative symptoms, while thalamic functional hypoconnectivity with multiple brain regions were significantly associated with specific negative symptom domains. rsFC of all 9 thalamic subregions involved some component of negative symptoms, and these thalamic hypoconnectivities with the temporoparietal junction-superior temporal sulcus (TPJ-STS), cerebellum, and striatum were associated with overall negative symptoms and some of its subdomains. These findings further raise the possibility that the thalamus may contribute to negative symptoms through its role in supporting communication and integration across distributed brain networks.

In this analysis, the thalamus was the only region that demonstrated both structural abnormalities in SSD patients relative to healthy controls and significant associations with BNSS-measured negative symptom severity. The thalamus has consistently emerged as a key locus of brain abnormalities in SSD, with large-scale structural and meta-analytic studies reporting robust reductions in thalamic volume among affected individuals ^15,16,73,74^, and a recent study suggests that abnormal thalamocortical development could contribute to emergence of SSD symptoms^75^. Several studies have also linked thalamic structural and functional abnormalities to negative symptom severity ^21,22,31,76^. However, the association was not necessary consistent ^20,21,77,78^. These inconsistencies may reflect differences in sample size, study populations, imaging methodologies, and symptom assessment approaches. The effect size of our structural findings compares favorably to other studies. For example, in the largest imaging study of SSD by ENIGMA Schizophrenia the SSD vs HC group differences had Cohen’s d =- 0.31, explaining about 2.3% of variance ^15^. Here, the Cohen’s d was -0.43 and associations with BNSS were ß =- 0.18, accounted for 4.6% of the variance, highlighting replicability and the likely clinically and statistically meaningful relationships. Guided by the structural findings, we subsequently focused on functional connectivity analyses on the thalamus. One of the strengths of this study is the use of prior thalamic subdivision method based on thalamic functional connectivity with major cortical networks. Given its central role in integrating and coordinating communication across multiple functional networks^50–54^, as well as accumulating evidence implicating large-scale network dysfunction in SSD and negative symptoms^22,23^, we believe that the adoption of this thalamic subdivision approach may provide important insights into negative symptom-related thalamic circuitry as compared to conventional histologically defined thalamic subdivisions. Although there is no 1:1 comparison with histological thalamic nuclei ^52^, some of the findings appear to be consistent with traditional understanding of histology-based thalamic nuclei. In contrast to prior studies which have reported both hyperconnectivity and hypoconnectivity within and between large-scale networks^22,23,39–49^, our findings were characterized predominantly by thalamic hypoconnectivity. This pattern is consistent with disrupted thalamic network integration in SSD and raises the possibility that thalamic dysfunction contributes to negative symptoms through impaired thalamic communication across neural systems.

More specifically, the rsFC of the somatomotor and medial temporal lobe related thalamic subregions with the TPJ-STS were significantly associated with overall negative symptom as well as blunted affect, and the reduced rsFC between medial temporal lobe-related thalamic subregion and TPJ-STS was also associated with anhedonia. Blunted affect was also associated with reduced rsFC between frontopariaetal, lateral occipital, and temporal thalamic subregions with TPJ-STS. Referencing histological thalamic nuclei, a majority of the medial temporal and part of somatomotor cortex-related thalamic subregions are consistent with medial pulvinar (PuM), which is known to have input from superior temporal cortex and function as part of the related associative network. In SSD population, disproportionally smaller PuM volume has been consistently shown ^79^, thought to be due to disrupted neurogenesis ^80^. Interestingly, we identified thalamocortical dysconnectivity only between several above-mentioned thalamic subregions and TPJ-STS, across negative symptoms sub-domains. The TPJ-STS area has been implicated in key human social cognitive functions: TPJ and posterior STS being responsible for theory of mind, complex semantic processing, social interaction, and anterior STS for sociality and emotional valence, with tendency of language-based information processing on the left and non-verbal processing on the right ^81–88^ and its dysfunction has been associated with negative symptoms in schizophrenia ^89–92^.

Reduced rsFC between default mode and medial occipital lobe-related thalamic subregions and cerebellum, specifically Crus I and Crus II, the cerebellar subregions implicated in theory of mind, mentalization, and self-related emotional processing ^93,94^, were associated with overall negative symptoms and blunted affect. In addition to these two subregions, executive and temporal thalamic subregions (equivalent to roughly the histological PuM and medial genicular nucleus) showed hypoconnectivities with the similar area of cerebellum associated with blunted affect. This finding supports previous studies identified link between emotion processing, social function, and motivation and the histological thalamic nuclei overlapping with default mode network-derived thalamic subregion, including mediodorsal nucleus ^95–97^, PulM^98–100^, and rostral intralaminar nucleus ^76,101^. Medial occipital lobe-related thalamic subregion mostly consists with inferior/lateral pulvinar, which is also associated with face-emotion, and social recognition^99,102,103^. The cerebellum, beyond its established role in motor coordination, contributes to a range of higher-order cognitive processes, including social cognition, ^104–107^ and have been linked to the development in psychopathology including psychosis^108–111^. Consistent with our findings, prior studies have linked cerebellar resting-state connectivity to social functioning^112,113^ as well as negative symptoms^25,27,114,115^, and notably, cerebellar TMS studies has shown changes in negative and cognitive symptoms and rsFC ^24,116^.

Blunted affect was also associated with hypoconnectivity between default mode, salience, somatomotor, and executive network-derived thalamic subregions and the striatum, including nucleus accumbens (NAcc). As a central component of mesolimbic dopaminergic circuitry, the NAcc has long been implicated in reward processing and reinforcement learning, and our findings are consistent with reported abnormality in NAcc associated with negative symptoms in SSD ^117–120^. Negative symptoms in SSD are also associated with striatal dopamine synthesis dysfunction and hypoconnectivity with mediodorsal thalamus ^27,121–123^, which is largely represented by default mode and salience thalamic subregions.

Several methodological strengths and limitations should be considered when interpreting these findings. First, this study employed the BNSS, which was developed to reflect the contemporary conceptualization of negative symptoms and permits symptom domain-specific assessment, in contrast to commonly used measures such as the SANS and PANSS that do not fully capture the current negative symptom framework ^124,125,12,22^. We believed this is also the first study on negative symptoms where the candidate regions were first empirically identified as exhibiting both structural alterations in SSD and associations with negative symptom severity, and subsequently used these findings to guide rsFC analyses, rather than relying solely on a priori region of interests or seeds. The thalamus was the only region meeting both criteria, providing data-driven support for focusing on thalamic networks. Furthermore, by using functionally defined thalamic subregions, we were able to assess connectivity across distinct thalamocortical, thalamosubcortical, and thalamocerebellar circuits, which may offer greater sensitivity to symptom-related network abnormalities than analyses treating the thalamus as a single structure.

The study has several limitations. Although negative symptom severity was associated with thalamic structural and functional abnormalities, the cross-sectional design is unable to prove causality. Replication in independent cohorts and longitudinal or interventional studies will be helpful to establish the robustness of these findings^126^ and for determining whether the thalamus functions as a causal node in negative symptoms. The SSD sample included individuals across a broad range of illness durations (average 16.4 years, SD 14.0), raising the possibility that some observed symptoms reflected secondary negative symptoms coming from overall disease severity, disease-related chronicity and related issues, or antipsychotic use ^12,13^. We found no significant association between negative symptoms and CPZ equivalent dose or positive symptoms, suggesting that the primary results may not be primarily driven by antipsychotic use. However, the contribution of other sources of secondary negative symptoms or antipsychotics’ effect on neuroimaging outcomes^127–133^ could not be fully ignored. Due to the voluntary nature of the clinical study involving MRI, there is a risk of selection bias where individuals with more severe negative symptoms might be less likely to participate. In addition, measurement of negative symptoms are currently depending on subjective report of internal and external experiences and raters subjective observation. Future study could be benefited from task-and ecological momentary assessment-based measurements for the validity. Furthermore, no significant rsFC associated with distress, asociality, avolition, or alogia were identified in this study, which might be the false negative findings although the sample size in this study is reasonable. Overall, extending the study to other non-thalamic related networks, or an even larger sample to identify rsFC linked to the negative symptom subdomains will be the follow up research step.

In conclusion, functional hypoconnectivity of thalamic regions linked to multiple functional networks were found to be associated with the overall negative symptoms in SSD. In addition, blunted affect was associated with the most extensive thalamic resting state functional connectivity abnormalities, while overlapping patterns of thalamic hypoconnectivity with TPJ-STS were observed in overall negative symptoms and anhedonia. These data support a potential role of thalamic-related network dysregulation in the overall negative symptoms and some of its subdomain symptoms. Future studies are implicated to clarify the causal relationships among these networks and to evaluate if targeted interventions to these thalamic networks that may improve negative symptoms.

## References

1. Foussias G, Agid O, Fervaha G, Remington G. Negative symptoms of schizophrenia: Clinical features, relevance to real world functioning and specificity versus other CNS disorders. Eur Neuropsychopharmacol. 2014;24(5):693–709. doi:10.1016/j.euroneuro.2013.10.017

2. Fusar-Poli P, Papanastasiou E, Stahl D, et al. Treatments of Negative Symptoms in Schizophrenia: Meta-Analysis of 168 Randomized Placebo-Controlled Trials. Schizophr Bull. 2015;41(4):892–899. doi:10.1093/schbul/sbu170

3. Galderisi S, Kaiser S, Bitter I, et al. EPA guidance on treatment of negative symptoms in schizophrenia. Eur Psychiatry. 2021;64(1):e21. doi:10.1192/j.eurpsy.2021.13

4. Németh G, Laszlovszky I, Czobor P, et al. Cariprazine versus risperidone monotherapy for treatment of predominant negative symptoms in patients with schizophrenia: a randomised, double-blind, controlled trial. Lancet. 2017;389(10074):1103–1113. doi:10.1016/S0140-6736(17)30060-0

5. Schneider-Thoma J, Zhu Y, Qin M, et al. Comparative efficacy and tolerability of antidopaminergic and muscarinic antipsychotics for acute schizophrenia: a network meta-analysis of randomised controlled trials indexed in international English and Chinese databases. Lancet. 2026;407(10531):876–891. doi:10.1016/S0140-6736(25)02365-7

6. Kaul I, Sawchak S, Correll CU, et al. Efficacy and safety of the muscarinic receptor agonist KarXT (xanomeline-trospium) in schizophrenia (EMERGENT-2) in the USA: results from a randomised, double-blind, placebo-controlled, flexible-dose phase 3 trial. Lancet. 2024;403(10422):160–170. doi:10.1016/S0140-6736(23)02190-6

7. Mizuno Y, McCutcheon RA, Brugger SP, Howes OD. Heterogeneity and efficacy of antipsychotic treatment for schizophrenia with or without treatment resistance: a meta-analysis. Neuropsychopharmacol Off Publ Am Coll Neuropsychopharmacol. 2020;45(4):622–631. doi:10.1038/s41386-019-0577-3

8. Galling B, Vernon JA, Pagsberg AK, et al. Efficacy and safety of antidepressant augmentation of continued antipsychotic treatment in patients with schizophrenia. Acta Psychiatr Scand. 2018;137(3):187–205. doi:10.1111/acps.12854

9. Helfer B, Samara MT, Huhn M, et al. Efficacy and Safety of Antidepressants Added to Antipsychotics for Schizophrenia: A Systematic Review and Meta-Analysis. Am J Psychiatry. 2016;173(9):876–886. doi:10.1176/appi.ajp.2016.15081035

10. Damiani S, Stefanelli R, Fortea L, et al. Interventions for negative symptoms in schizophrenia: efficacy and clinical interpretability in a meta-analysis of 451 randomized controlled trials. Mol Psychiatry. 2026;31(8):4259–4269. doi:10.1038/s41380-026-03543-1

11. Buchanan RW, Katz I, Issa F, et al. Pharmacological and Somatic Treatments for First-Episode Psychosis and Schizophrenia: Synopsis of the US Department of Veterans Affairs and US Department of Defense Clinical Practice Guidelines. Schizophr Bull. 2025;51(4):969–982. doi:10.1093/schbul/sbaf053

12. Correll CU, Schooler NR. Negative Symptoms in Schizophrenia: A Review and Clinical Guide for Recognition, Assessment, and Treatment. Neuropsychiatr Dis Treat. 2020;16:519–534. doi:10.2147/NDT.S225643

13. Govil P, Kantrowitz JT. Negative Symptoms in Schizophrenia: An Update on Research Assessment and the Current and Upcoming Treatment Landscape. CNS Drugs. 2025;39(3):243–262. doi:10.1007/s40263-024-01151-7

14. van Erp TGM, Walton E, Hibar DP, et al. Cortical Brain Abnormalities in 4474 Individuals With Schizophrenia and 5098 Control Subjects via the Enhancing Neuro Imaging Genetics Through Meta Analysis (ENIGMA) Consortium. Biol Psychiatry. 2018;84(9):644–654. doi:10.1016/j.biopsych.2018.04.023

15. van Erp TGM, Hibar DP, Rasmussen JM, et al. Subcortical brain volume abnormalities in 2028 individuals with schizophrenia and 2540 healthy controls via the ENIGMA consortium. Mol Psychiatry. 2016;21(4):547–553. doi:10.1038/mp.2015.63

16. Lawrie SM. Brain structure in schizophrenia: what do we know and what next? Lancet Psychiatry. 2025;12(11):852–862. doi:10.1016/S2215-0366(25)00172-5

17. Banaj N, Vecchio D, Piras F, et al. Cortical morphology in patients with the deficit and non-deficit syndrome of schizophrenia: a worldwide meta- and mega-analyses. Mol Psychiatry. 2023;28(10):4363–4373. doi:10.1038/s41380-023-02221-w

18. Kamalakannan SMV, Male AG, Yilanli M, et al. Neuroanatomical Correlates of Negative Symptoms in Schizophrenia. bioRxiv. Preprint posted online January 25, 2026:2025.09.22.677864. doi:10.1101/2025.09.22.677864

19. Walton E, Hibar DP, Erp TGM van, et al. Prefrontal cortical thinning links to negative symptoms in schizophrenia via the ENIGMA consortium. Psychol Med. 2018;48(1):82–94. doi:10.1017/S0033291717001283

20. Zhou C, Zhang R, Ding M, et al. Progressive structural alterations associated with negative symptoms in schizophrenia: A causal structural covariance network analysis. Prog Neuropsychopharmacol Biol Psychiatry. 2025;136:111236. doi:10.1016/j.pnpbp.2024.111236

21. Zhu T, Wang Z, Zhou C, et al. Meta-analysis of structural and functional brain abnormalities in schizophrenia with persistent negative symptoms using activation likelihood estimation. Front Psychiatry. 2022;13:957685. doi:10.3389/fpsyt.2022.957685

22. Dalloul N, van Huijgevoort C, Belmont J, Barch DM. Negative symptoms and resting-state functional connectivity across psychotic and mood disorders: A systematic literature review. Neurosci Biobehav Rev. 2025;178:106385. doi:10.1016/j.neubiorev.2025.106385

23. Dong D, Wang Y, Chang X, Luo C, Yao D. Dysfunction of Large-Scale Brain Networks in Schizophrenia: A Meta-analysis of Resting-State Functional Connectivity. Schizophr Bull. 2018;44(1):168–181. doi:10.1093/schbul/sbx034

24. Brady RO, Gonsalvez I, Lee I, et al. Cerebellar-Prefrontal Network Connectivity and Negative Symptoms in Schizophrenia. Am J Psychiatry. 2019;176(7):512–520. doi:10.1176/appi.ajp.2018.18040429

25. Choi SY, Ha M, Choi S, et al. Altered intrinsic cerebellar-cerebral functional connectivity is related to negative symptoms in patients with first-episode psychosis. Schizophr Res. 2023;252:56–63. doi:10.1016/j.schres.2022.12.041

26. Li Z, Lei W, Deng W, et al. Aberrant spontaneous neural activity and correlation with evoked-brain potentials in first-episode, treatment-naïve patients with deficit and non-deficit schizophrenia. Psychiatry Res Neuroimaging. 2017;261:9–19. doi:10.1016/j.pscychresns.2017.01.001

27. Wang X, Guo X dong, Cai X lu, et al. Cerebellar–Basal Ganglia Dysconnectivity in Schizophrenia: Insights into Motivational Deficits. Schizophr Bull. 2026;52(2):sbaf087. doi:10.1093/schbul/sbaf087

28. Fang J, Cai R, Hu Y, et al. Aberrant brain functional connectivity mediates the effects of negative symptoms on cognitive function in schizophrenia: A structural equation model. J Psychiatr Res. 2024;177:109–117. doi:10.1016/j.jpsychires.2024.07.006

29. Forlim CG, Klock L, Bächle J, et al. Reduced Resting-State Connectivity in the Precuneus is correlated with Apathy in Patients with Schizophrenia. Sci Rep. 2020;10(1):2616. doi:10.1038/s41598-020-59393-6

30. Zhu T, Wang Z, Wu W, et al. Altered brain functional networks in schizophrenia with persistent negative symptoms: an activation likelihood estimation meta-analysis. Front Hum Neurosci. 2023;17:1204632. doi:10.3389/fnhum.2023.1204632

31. Brandl F, Avram M, Weise B, et al. Specific Substantial Dysconnectivity in Schizophrenia: A Transdiagnostic Multimodal Meta-analysis of Resting-State Functional and Structural Magnetic Resonance Imaging Studies. Biol Psychiatry. 2019;85(7):573–583. doi:10.1016/j.biopsych.2018.12.003

32. Giordano GM, Stanziano M, Papa M, et al. Functional connectivity of the ventral tegmental area and avolition in subjects with schizophrenia: a resting state functional MRI study. Eur Neuropsychopharmacol J Eur Coll Neuropsychopharmacol. 2018;28(5):589–602. doi:10.1016/j.euroneuro.2018.03.013

33. Xu P, Klaasen NG, Opmeer EM, et al. Intrinsic mesocorticolimbic connectivity is negatively associated with social amotivation in people with schizophrenia. Schizophr Res. 2019;208:353–359. doi:10.1016/j.schres.2019.01.023

34. Brakowski J, Manoliu A, Homan P, et al. Aberrant striatal coupling with default mode and central executive network relates to self-reported avolition and anhedonia in schizophrenia. J Psychiatr Res. 2022;145:263–275. doi:10.1016/j.jpsychires.2020.10.047

35. Geffen T, Hardikar S, Smallwood J, et al. Striatal Functional Hypoconnectivity in Patients With Schizophrenia Suffering From Negative Symptoms, Longitudinal Findings. Schizophr Bull. 2024;50(6):1337–1348. doi:10.1093/schbul/sbae052

36. Nawaz U, Lee I, Beermann A, Eack S, Keshavan M, Brady R. Individual Variation in Functional Brain Network Topography is Linked to Schizophrenia Symptomatology. Schizophr Bull. 2021;47(1):180–188. doi:10.1093/schbul/sbaa088

37. Shukla DK, Chiappelli JJ, Sampath H, et al. Aberrant Frontostriatal Connectivity in Negative Symptoms of Schizophrenia. Schizophr Bull. 2019;45(5):1051–1059. doi:10.1093/schbul/sby165

38. Zhang M, Zheng Y, Zhang J, et al. Decoupling of amygdala-salience circuit associated with negative symptoms in adolescents with early-onset schizophrenia. Psychiatry Clin Neurosci. Published online May 2026. doi:10.1111/pcn.70080

39. Biondi M, Marino M, Mantini D, Spironelli C. Unveiling altered connectivity between cognitive networks and cerebellum in schizophrenia. Schizophr Res. 2024;271:47–58. doi:10.1016/j.schres.2024.06.044

40. Gallucci J, Yu JC, Oliver LD, et al. Neural Circuitry and Therapeutic Targeting of Depressive Symptoms in Schizophrenia Spectrum Disorders. Am J Psychiatry. 2024;181:910–919. doi:10.1176/appi.ajp.20230655

41. Kebets V, Holmes AJ, Orban C, et al. Somatosensory-Motor Dysconnectivity Spans Multiple Transdiagnostic Dimensions of Psychopathology. Biol Psychiatry. 2019;86(10):779–791. doi:10.1016/j.biopsych.2019.06.013

42. Lee WH, Doucet GE, Leibu E, Frangou S. Resting-state network connectivity and metastability predict clinical symptoms in schizophrenia. Schizophr Res. 2018;201:208–216. doi:10.1016/j.schres.2018.04.029

43. Li T, Wang Q, Zhang J, et al. Brain-Wide Analysis of Functional Connectivity in First-Episode and Chronic Stages of Schizophrenia. Schizophr Bull. 2017;43(2):436–448. doi:10.1093/schbul/sbw099

44. Mellem MS, Liu Y, Gonzalez H, Kollada M, Martin WJ, Ahammad P. Machine Learning Models Identify Multimodal Measurements Highly Predictive of Transdiagnostic Symptom Severity for Mood, Anhedonia, and Anxiety. Biol Psychiatry Cogn Neurosci Neuroimaging. 2020;5(1):56–67. doi:10.1016/j.bpsc.2019.07.007

45. Wang D, Li M, Wang M, et al. Individual-specific functional connectivity markers track dimensional and categorical features of psychotic illness. Mol Psychiatry. 2020;25(9):2119–2129. doi:10.1038/s41380-018-0276-1

46. Xi YB, Guo F, Liu WM, et al. Triple network hypothesis-related disrupted connections in schizophrenia: A spectral dynamic causal modeling analysis with functional magnetic resonance imaging. Schizophr Res. 2021;233:89–96. doi:10.1016/j.schres.2021.06.024

47. Zapetis SL, Nasiriavanaki Z, Luther L, Holt DJ. Neural Correlates of Variation in Personal Space and Social Functioning in Schizophrenia and Healthy Individuals. Schizophr Bull. 2022;48(5):1075–1084. doi:10.1093/schbul/sbac052

48. Zhang Y jing, Pu C cheng, Wang Y ming, et al. Social brain network correlates with real-life social network in individuals with schizophrenia and social anhedonia. Schizophr Res. 2021;232:77–84. doi:10.1016/j.schres.2021.05.016

49. Zhao C, Jiang R, Bustillo J, et al. Cross-cohort replicable resting-state functional connectivity in predicting symptoms and cognition of schizophrenia. Hum Brain Mapp. 2024;45(7):e26694. doi:10.1002/hbm.26694

50. Greene DJ, Marek S, Gordon EM, et al. Integrative and Network-Specific Connectivity of the Basal Ganglia and Thalamus Defined in Individuals. Neuron. 2020;105(4):742–758.e6. doi:10.1016/j.neuron.2019.11.012

51. Halassa MM, Kastner S. Thalamic functions in distributed cognitive control. Nat Neurosci. 2017;20(12):1669–1679. doi:10.1038/s41593-017-0020-1

52. Hwang K, Bertolero MA, Liu WB, D’Esposito M. The Human Thalamus Is an Integrative Hub for Functional Brain Networks. J Neurosci Off J Soc Neurosci. 2017;37(23):5594–5607. doi:10.1523/JNEUROSCI.0067-17.2017

53. Hwang K, Shine JM, Bruss J, Tranel D, Boes A. Neuropsychological evidence of multi-domain network hubs in the human thalamus. Diedrichsen J, Behrens TE, Dosenbach N, eds. eLife. 2021;10:e69480. doi:10.7554/eLife.69480

54. Yuan R, Di X, Taylor PA, Gohel S, Tsai YH, Biswal BB. Functional topography of the thalamocortical system in human. Brain Struct Funct. 2016;221(4):1971–1984. doi:10.1007/s00429-015-1018-7

55. Gardner DM, Murphy AL, O’Donnell H, Centorrino F, Baldessarini RJ. International consensus study of antipsychotic dosing. Am J Psychiatry. 2010;167(6):686–693. doi:10.1176/appi.ajp.2009.09060802

56. McAdam MK, Baldessarini RJ, Murphy AL, Gardner DM. Second International Consensus Study of Antipsychotic Dosing (ICSAD-2). J Psychopharmacol Oxf Engl. 2023;37(10):982–991. doi:10.1177/02698811231205688

57. Kirkpatrick B, Strauss GP, Nguyen L, et al. The brief negative symptom scale: psychometric properties. Schizophr Bull. 2011;37(2):300–305. doi:10.1093/schbul/sbq059

58. Shafer A. Meta-analysis of the Brief Psychiatric Rating Scale factor structure. Psychol Assess. 2005;17(3):324–335. doi:10.1037/1040-3590.17.3.324

59. Wechsler Adult Intelligence Scale--Third Edition. Accessed October 3, 2026. https://psycnet.apa.org/doiLanding?doi=10.1037%2Ft49755-000

60. Keefe RSE, Goldberg TE, Harvey PD, Gold JM, Poe MP, Coughenour L. The Brief Assessment of Cognition in Schizophrenia: reliability, sensitivity, and comparison with a standard neurocognitive battery. Schizophr Res. 2004;68(2-3):283–297. doi:10.1016/j.schres.2003.09.011

61. Fischl B. FreeSurfer. NeuroImage. 2012;62(2):774–781. doi:10.1016/j.neuroimage.2012.01.021

62. Desikan RS, Ségonne F, Fischl B, et al. An automated labeling system for subdividing the human cerebral cortex on MRI scans into gyral based regions of interest. NeuroImage. 2006;31(3):968–980. doi:10.1016/j.neuroimage.2006.01.021

63. Fischl B, Salat DH, Busa E, et al. Whole brain segmentation: automated labeling of neuroanatomical structures in the human brain. Neuron. 2002;33(3):341–355. doi:10.1016/s0896-6273(02)00569-x

64. Fortin JP, Cullen N, Sheline YI, et al. Harmonization of cortical thickness measurements across scanners and sites. NeuroImage. 2018;167:104–120. doi:10.1016/j.neuroimage.2017.11.024

65. Cox RW. AFNI: software for analysis and visualization of functional magnetic resonance neuroimages. Comput Biomed Res Int J. 1996;29(3):162–173. doi:10.1006/cbmr.1996.0014

66. Adhikari BM, Jahanshad N, Shukla D, et al. Heritability estimates on resting state fMRI data using ENIGMA analysis pipeline. Pac Symp Biocomput Pac Symp Biocomput. 2018;23:307–318.

67. Adhikari BM, Jahanshad N, Shukla D, et al. A resting state fMRI analysis pipeline for pooling inference across diverse cohorts:An ENIGMA rs-fMRI Protocol. Brain Imaging Behav. 2019;13(5):1453–1467. doi:10.1007/s11682-018-9941-x

68. Dickie EW, Shahab S, Hawco C, et al. Robust hierarchically organized whole-brain patterns of dysconnectivity in schizophrenia spectrum disorders observed after personalized intrinsic network topography. Hum Brain Mapp. 2023;44(15):5153–5166. doi:10.1002/hbm.26453

69. Du X, Wei X, Ding H, et al. Unraveling schizophrenia replicable functional connectivity disruption patterns across sites. Hum Brain Mapp. 2023;44(1):156–169. doi:10.1002/hbm.26108

70. Butry L, Thomä J, Forsting J, Enax-Krumova EK, Schlaffke L. Cross-vendor reliability of functional and structural brain connectivity in a travelling cohort. Sci Rep. 2026;16(1):12071. doi:10.1038/s41598-026-47705-1

71. Yamashita A, Yahata N, Itahashi T, et al. Harmonization of resting-state functional MRI data across multiple imaging sites via the separation of site differences into sampling bias and measurement bias. PLOS Biol. 2019;17(4):e3000042. doi:10.1371/journal.pbio.3000042

72. Yu M, Linn KA, Cook PA, et al. Statistical harmonization corrects site effects in functional connectivity measurements from multi-site fMRI data. Hum Brain Mapp. 2018;39(11):4213–4227. doi:10.1002/hbm.24241

73. Cho KIK, Shenton ME, Kubicki M, et al. Altered Thalamo-Cortical White Matter Connectivity: Probabilistic Tractography Study in Clinical-High Risk for Psychosis and First-Episode Psychosis. Schizophr Bull. 2016;42(3):723–731. doi:10.1093/schbul/sbv169

74. Kubota M, Miyata J, Sasamoto A, et al. Thalamocortical Disconnection in the Orbitofrontal Region Associated With Cortical Thinning in Schizophrenia. JAMA Psychiatry. 2013;70(1):12–21. doi:10.1001/archgenpsychiatry.2012.1023

75. Benoit LJ, Canetta S, Kellendonk C. Thalamocortical Development: A Neurodevelopmental Framework for Schizophrenia. Biol Psychiatry. 2022;92(6):491–500. doi:10.1016/j.biopsych.2022.03.004

76. Bayrakçı A, Zorlu N, Karakılıç M, et al. Negative symptoms are associated with modularity and thalamic connectivity in schizophrenia. Eur Arch Psychiatry Clin Neurosci. 2023;273(3):565–574. doi:10.1007/s00406-022-01433-5

77. Ferri J, Ford JM, Roach BJ, et al. Resting-state thalamic dysconnectivity in schizophrenia and relationships with symptoms. Psychol Med. 2018;48(15):2492–2499. doi:10.1017/S003329171800003X

78. Kirschner M, Schmidt A, Hodzic-Santor B, et al. Orbitofrontal-Striatal Structural Alterations Linked to Negative Symptoms at Different Stages of the Schizophrenia Spectrum. Schizophr Bull. 2021;47(3):849–863. doi:10.1093/schbul/sbaa169

79. Dorph-Petersen KA, Lewis DA. Postmortem structural studies of the thalamus in schizophrenia. Schizophr Res. 2017;180:28–35. doi:10.1016/j.schres.2016.08.007

80. Byne W, Fernandes J, Haroutunian V, et al. Reduction of right medial pulvinar volume and neuron number in schizophrenia. Schizophr Res. 2007;90(1-3):71–75. doi:10.1016/j.schres.2006.10.006

81. Deen B, Koldewyn K, Kanwisher N, Saxe R. Functional Organization of Social Perception and Cognition in the Superior Temporal Sulcus. Cereb Cortex. 2015;25(11):4596–4609. doi:10.1093/cercor/bhv111

82. Henry JD, von Hippel W, Molenberghs P, Lee T, Sachdev PS. Clinical assessment of social cognitive function in neurological disorders. Nat Rev Neurol. 2016;12(1):28–39. doi:10.1038/nrneurol.2015.229

83. Jabbi M, Kohn PD, Nash T, et al. Convergent BOLD and Beta-Band Activity in Superior Temporal Sulcus and Frontolimbic Circuitry Underpins Human Emotion Cognition. Cereb Cortex. 2015;25(7):1878–1888. doi:10.1093/cercor/bht427

84. McMahon E, Isik L. Seeing Social Interactions. Trends Cogn Sci. 2023;27(12):1165–1179. doi:10.1016/j.tics.2023.09.001

85. Patel GH, Sestieri C, Corbetta M. The evolution of the temporoparietal junction and posterior superior temporal sulcus. Cortex J Devoted Study Nerv Syst Behav. 2019;118:38–50. doi:10.1016/j.cortex.2019.01.026

86. Pobric G, Lambon Ralph MA, Zahn R. Hemispheric Specialization within the Superior Anterior Temporal Cortex for Social and Nonsocial Concepts. J Cogn Neurosci. 2016;28(3):351–360. doi:10.1162/jocn_a_00902

87. Sato W, Kochiyama T, Uono S, et al. Widespread and lateralized social brain activity for processing dynamic facial expressions. Hum Brain Mapp. 2019;40(13):3753–3768. doi:10.1002/hbm.24629

88. Wang X, Wang B, Bi Y. Close yet independent: Dissociation of social from valence and abstract semantic dimensions in the left anterior temporal lobe. Hum Brain Mapp. 2019;40(16):4759–4776. doi:10.1002/hbm.24735

89. Bodnar M, Hovington CL, Buchy L, Malla AK, Joober R, Lepage M. Cortical thinning in temporo-parietal junction (TPJ) in non-affective first-episode of psychosis patients with persistent negative symptoms. PloS One. 2014;9(6):e101372. doi:10.1371/journal.pone.0101372

90. Demjaha A, Galderisi S, Glenthøj B, et al. Negative symptoms in First-Episode Schizophrenia related to morphometric alterations in orbitofrontal and superior temporal cortex: the OPTiMiSE study. Psychol Med. 2023;53(8):3471–3479. doi:10.1017/S0033291722000010

91. Gao J, Jiang R, Tang X, et al. A neuromarker for deficit syndrome in schizophrenia from a combination of structural and functional magnetic resonance imaging. CNS Neurosci Ther. 2023;29(12):3774–3785. doi:10.1111/cns.14297

92. Patel GH, Arkin SC, Ruiz-Betancourt DR, et al. Failure to engage the temporoparietal junction/posterior superior temporal sulcus predicts impaired naturalistic social cognition in schizophrenia. Brain. 2021;144(6):1898–1910. doi:10.1093/brain/awab081

93. Beuriat PA, Cohen-Zimerman S, Smith GNL, Krueger F, Gordon B, Grafman J. Evidence of the role of the cerebellum in cognitive theory of mind using voxel-based lesion mapping. Sci Rep. 2022;12(1):4999. doi:10.1038/s41598-022-09104-0

94. Van Overwalle F, Ma Q, Heleven E. The posterior crus II cerebellum is specialized for social mentalizing and emotional self-experiences: a meta-analysis. Soc Cogn Affect Neurosci. 2020;15(9):905–928. doi:10.1093/scan/nsaa124

95. Golden EC, Graff-Radford J, Jones DT, Benarroch EE. Mediodorsal nucleus and its multiple cognitive functions. Neurology. 2016;87(20):2161–2168. doi:10.1212/WNL.0000000000003344

96. Jakary A, Vinogradov S, Feiwell R, Deicken RF. N-acetylaspartate reductions in the mediodorsal and anterior thalamus in men with schizophrenia verified by tissue volume corrected proton MRSI. Schizophr Res. 2005;76(2-3):173–185. doi:10.1016/j.schres.2005.02.012

97. Mørch-Johnsen L, Jørgensen KN, Barth C, et al. Thalamic nuclei volumes in schizophrenia and bipolar spectrum disorders - Associations with diagnosis and clinical characteristics. Schizophr Res. 2023;256:26–35. doi:10.1016/j.schres.2023.04.008

98. Alemán-Gómez Y, Baumgartner T, Klauser P, et al. Multimodal Magnetic Resonance Imaging Depicts Widespread and Subregion Specific Anomalies in the Thalamus of Early-Psychosis and Chronic Schizophrenia Patients. Schizophr Bull. 2023;49(1):196–207. doi:10.1093/schbul/sbac113

99. Martínez A, Tobe RH, Gaspar PA, et al. Disease-Specific Contribution of Pulvinar Dysfunction to Impaired Emotion Recognition in Schizophrenia. Front Behav Neurosci. 2021;15:787383. doi:10.3389/fnbeh.2021.787383

100. Zhang Y, Lu Z, Sun Y, et al. Predictive role of pulvinar in social functional outcome of schizophrenia. Psychiatry Res. 2023;327:115419. doi:10.1016/j.psychres.2023.115419

101. Vertes RP, Linley SB, Rojas AKP. Structural and functional organization of the midline and intralaminar nuclei of the thalamus. Front Behav Neurosci. 2022;16:964644. doi:10.3389/fnbeh.2022.964644

102. Cortes N, Mandel C, Ikan L, Casanova C. The pulvinar nucleus and its role in cognitive functions. Brain Res Bull. 2026;234:111672. doi:10.1016/j.brainresbull.2025.111672

103. Martínez A, Gaspar PA, Hillyard SA, et al. Impaired Motion Processing in Schizophrenia and the Attenuated Psychosis Syndrome: Etiological and Clinical Implications. Am J Psychiatry. 2018;175(12):1243–1254. doi:10.1176/appi.ajp.2018.18010072

104. King M, Hernandez-Castillo CR, Poldrack RA, Ivry RB, Diedrichsen J. Functional boundaries in the human cerebellum revealed by a multi-domain task battery. Nat Neurosci. 2019;22(8):1371–1378. doi:10.1038/s41593-019-0436-x

105. Metoki A, Wang Y, Olson IR. The Social Cerebellum: A Large-Scale Investigation of Functional and Structural Specificity and Connectivity. Cereb Cortex. 2022;32(5):987–1003. doi:10.1093/cercor/bhab260

106. Nettekoven C, Zhi D, Shahshahani L, et al. A hierarchical atlas of the human cerebellum for functional precision mapping. Nat Commun. 2024;15(1):8376. doi:10.1038/s41467-024-52371-w

107. Van Overwalle F, Manto M, Cattaneo Z, et al. Consensus Paper: Cerebellum and Social Cognition. Cerebellum. 2020;19(6):833–868. doi:10.1007/s12311-020-01155-1

108. Andreasen NC, Pierson R. The Role of the Cerebellum in Schizophrenia. Biol Psychiatry. 2008;64(2):81–88. doi:10.1016/j.biopsych.2008.01.003

109. Hughes DE, Kunitoki K, Elyounssi S, et al. Genetic patterning for child psychopathology is distinct from that for adults and implicates fetal cerebellar development. Nat Neurosci. 2023;26(6):959–969. doi:10.1038/s41593-023-01321-8

110. Moussa-Tooks AB, Rogers BP, Huang AS, Sheffield JM, Heckers S, Woodward ND. CEREBELLAR STRUCTURE AND COGNITIVE ABILITY IN PSYCHOSIS. Biol Psychiatry. 2022;92(5):385–395. doi:10.1016/j.biopsych.2022.03.013

111. Sathyanesan A, Zhou J, Scafidi J, Heck DH, Sillitoe RV, Gallo V. Emerging connections between cerebellar development, behaviour and complex brain disorders. Nat Rev Neurosci. 2019;20(5):298–313. doi:10.1038/s41583-019-0152-2

112. Moe AM, Blain SD, Kalathil A, et al. Contributions of the posterior cerebellum to mentalizing and social functioning: A transdiagnostic investigation. Psychol Med. 2025;55:e67. doi:10.1017/S003329172500039X

113. Mothersill O, Knee-Zaska C, Donohoe G. Emotion and Theory of Mind in Schizophrenia-Investigating the Role of the Cerebellum. Cerebellum. 2016;15(3):357–368. doi:10.1007/s12311-015-0696-2

114. Awada J, Delavari F, Bolton TAW, et al. A Longitudinal and Reproducible Anti-Coactivation Pattern Between the Cerebellum and the Ventral Tegmental Area Is Related to Apathy in Schizophrenia. Biol Psychiatry. 2026;99(2):124–133. doi:10.1016/j.biopsych.2025.06.009

115. Cai XL, Wang YM, Wang Y, et al. Neurological Soft Signs Are Associated With Altered Cerebellar-Cerebral Functional Connectivity in Schizophrenia. Schizophr Bull. 2021;47(5):1452–1462. doi:10.1093/schbul/sbaa200

116. Cao H, Argyelan M, Yan J, et al. Mapping cerebellar connectivity to cognition in psychosis: Convergent evidence from fMRI and TMS. Biol Psychiatry. 2026;99(2):134–141. doi:10.1016/j.biopsych.2025.06.023

117. Juckel G, Schlagenhauf F, Koslowski M, et al. Dysfunction of ventral striatal reward prediction in schizophrenia. NeuroImage. 2006;29(2):409–416. doi:10.1016/j.neuroimage.2005.07.051

118. van Gool R, Cay M, van der Heijden H, et al. Neural and Behavioral Markers of Negative Symptoms and Cognitive Impairment in Patients with Psychosis. J Neurosci Off J Soc Neurosci. 2025;45(30):e0039252025. doi:10.1523/JNEUROSCI.0039-25.2025

119. Zeng J, Yan J, Cao H, et al. Neural substrates of reward anticipation and outcome in schizophrenia: a meta-analysis of fMRI findings in the monetary incentive delay task. Transl Psychiatry. 2022;12:448. doi:10.1038/s41398-022-02201-8

120. Zhou C, Xue C, Chen J, et al. Altered Functional Connectivity of the Nucleus Accumbens Network Between Deficit and Non-deficit Schizophrenia. Front Psychiatry. 2021;12:704631. doi:10.3389/fpsyt.2021.704631

121. Avram M, Brandl F, Bäuml J, Sorg C. Cortico-thalamic hypo- and hyperconnectivity extend consistently to basal ganglia in schizophrenia. Neuropsychopharmacology. 2018;43(11):2239–2248. doi:10.1038/s41386-018-0059-z

122. Avram M, Brandl F, Knolle F, et al. Aberrant striatal dopamine links topographically with cortico-thalamic dysconnectivity in schizophrenia. Brain J Neurol. 2020;143(11):3495–3505. doi:10.1093/brain/awaa296

123. Brandl F, Knolle F, Avram M, et al. Negative symptoms, striatal dopamine and model-free reward decision-making in schizophrenia. Brain J Neurol. 2023;146(2):767–777. doi:10.1093/brain/awac268

124. Kirkpatrick B, Fenton WS, Carpenter WT, Marder SR. The NIMH-MATRICS consensus statement on negative symptoms. Schizophr Bull. 2006;32(2):214–219. doi:10.1093/schbul/sbj053

125. Marder SR, Galderisi S. The current conceptualization of negative symptoms in schizophrenia. World Psychiatry Off J World Psychiatr Assoc. 2017;16(1):14–24. doi:10.1002/wps.20385

126. Siddiqi SH, Kording KP, Parvizi J, Fox MD. Causal mapping of human brain function. Nat Rev Neurosci. 2022;23(6):361–375. doi:10.1038/s41583-022-00583-8

127. Chopra S, Fornito A, Francey SM, et al. Differentiating the effect of antipsychotic medication and illness on brain volume reductions in first-episode psychosis: A Longitudinal, Randomised, Triple-blind, Placebo-controlled MRI Study. Neuropsychopharmacol Off Publ Am Coll Neuropsychopharmacol. 2021;46(8):1494–1501. doi:10.1038/s41386-021-00980-0

128. Chopra S, Francey SM, O’Donoghue B, et al. Functional Connectivity in Antipsychotic-Treated and Antipsychotic-Naive Patients With First-Episode Psychosis and Low Risk of Self-harm or Aggression: A Secondary Analysis of a Randomized Clinical Trial. JAMA Psychiatry. 2021;78(9):994–1004. doi:10.1001/jamapsychiatry.2021.1422

129. Fusar-Poli P, Smieskova R, Kempton MJ, Ho BC, Andreasen NC, Borgwardt S. Progressive brain changes in schizophrenia related to antipsychotic treatment? A meta-analysis of longitudinal MRI studies. Neurosci Biobehav Rev. 2013;37(8):1680–1691. doi:10.1016/j.neubiorev.2013.06.001

130. Jiang Y, Wang Y, Huang H, et al. Antipsychotics Effects on Network-Level Reconfiguration of Cortical Morphometry in First-Episode Schizophrenia. Schizophr Bull. 2022;48(1):231–240. doi:10.1093/schbul/sbab082

131. Lesh TA, Tanase C, Geib BR, et al. A Multimodal Analysis of Antipsychotic Effects on Brain Structure and Function in First-Episode Schizophrenia. JAMA Psychiatry. 2015;72(3):226–234. doi:10.1001/jamapsychiatry.2014.2178

132. Lui S, Li T, Deng W, et al. Short-term Effects of Antipsychotic Treatment on Cerebral Function in Drug-Naive First-Episode Schizophrenia Revealed by “Resting State” Functional Magnetic Resonance Imaging. Arch Gen Psychiatry. 2010;67(8):783–792. doi:10.1001/archgenpsychiatry.2010.84

133. Vita A, De Peri L, Deste G, Barlati S, Sacchetti E. The Effect of Antipsychotic Treatment on Cortical Gray Matter Changes in Schizophrenia: Does the Class Matter? A Meta-analysis and Meta-regression of Longitudinal Magnetic Resonance Imaging Studies. Biol Psychiatry. 2015;78(6):403–412. doi:10.1016/j.biopsych.2015.02.008

134. Jakab A, Blanc R, Szekely G. Morel Atlas of the Human Thalamus (MNI152 space, voxelized version). Published online October 2024. doi:10.5281/zenodo.13918589

135. Krauth A, Blanc R, Poveda A, Jeanmonod D, Morel A, Székely G. A mean three-dimensional atlas of the human thalamus: generation from multiple histological data. NeuroImage. 2010;49(3):2053–2062. doi:10.1016/j.neuroimage.2009.10.042

136. Morel A. Stereotactic Atlas of the Human Thalamus and Basal Ganglia. CRC Press; 2007. doi:10.3109/9781420016796

137. Morel A, Magnin M, Jeanmonod D. Multiarchitectonic and stereotactic atlas of the human thalamus. J Comp Neurol. 1997;387(4):588–630. doi:10.1002/(SICI)1096-9861(19971103)387:4<588::AID-CNE8>3.0.CO;2-Z

